# Energy Landscape near Glioma Decipher the Role of Glioma Stem Cells within Glioma Hierarchy

**DOI:** 10.1101/2023.11.24.568616

**Authors:** Mengchao Yao, Yang Su, Ruiqi Xiong, Xile Zhang, Xiao-Mei Zhu, Yong-Cong Chen, Ping Ao

## Abstract

Glioma stem cells (GSCs) have been recognized as key players in glioma recurrence and therapeutic resistance, presenting a promising target for novel treatments. However, the limited understanding of the role GSCs play in the glioma hierarchy has drawn controversy and hindered research translation into therapies. Despite significant advances in our understanding of gene regulatory networks, the dynamics of these networks and their implications for glioma remain elusive. This study employs a systemic theoretical perspective to integrate empirical knowledge into a core endogenous network model for glioma, thereby elucidating its energy landscape through network dynamics computation. The model identifies two stable states corresponding to astrocytoma and oligodendroglioma, connected by a transitional state associated with GSCs, indicating the instability of GSCs *in vivo* and providing negative proof of their role as the cellular origin for glioma. We also obtained various stable states further indicative of glioma’s multicellular origins and uncovered a group of transition states that could potentially induce tumor heterogeneity and therapeutic resistance. This research proposes that the transitional states linking both glioma stable states are central to glioma heterogeneity and therapy resistance, suggesting a combination of apoptosis-inducing and differentiation-promoting therapies as a potential strategy. Our approach holds promise for advancing glioma therapy by providing a new lens through which to view the complex landscape of glioma biology.

## Introduction

Years ago, glioma cells with the characteristics of neural/glia stem cells were identified as cancer stem cells in glioma or glioma stem cells (GSCs) (Hemmati et al. 2003; Ignatova et al. 2002; Galli et al. 2004). GSCs play crucial roles in the recurrence of glioma and the resistance to radiation and chemotherapy (Biserova et al. 2021; Lathia et al. 2015) and thus are considered promising therapeutic targets for glioma. The glioma hierarchy model was proposed, depicting all cells within the glioma as forming a Waddington epigenetic landscape-like structure, where GSCs are put at the apex (Lathia et al. 2015; Gimple et al. 2019). However, limited understanding of GSCs has drawn controversy (Rahman et al. 2011; Capp 2019) and hindered research translation into therapies (Lathia et al. 2015). Whether GSCs are the cells of glioma origin remains unclear (Biserova et al. 2021; Lathia et al. 2015; Gimple et al. 2019), the intermediate states between somatic glia and glioma cells are still hiding behind haze, even the views of GSCs and non-stem tumor cells (NSTCs) are mutually exclusive in the glioma hierarchy (Lathia et al. 2015).

Gliomas and the normal glial system may follow the same differentiation rules (Laug, Glasgow, and Deneen 2018), understanding the principle behind the carcinogenesis of glioma and the development process of the glial system, that is, the gene regulatory network and the landscape determined by it (Ao 2008; Huang, Ernberg, and Kauffman 2009; Raju and Siggia 2023; Feinberg and Levchenko 2023), may lead to the enlightenments of the role of GSCs in glioma hierarchy, thereby helping to find insights for new therapies (Lathia et al. 2015; Uthamacumaran 2022; Prokop 2022).

Currently, data-driven studies are attempting to obtain the gene regulatory network and landscape of glioma and the glial system based on high-throughput data (Csermely et al. 2015; Niu et al. 2019; Qiu et al. 2022; Feinberg and Levchenko 2023). Notably, Hoang *et al*. (Hoang et al. 2020) have driven gene regulatory networks of retinal regeneration from data, Suvà and Tirosh (Suvà and Tirosh 2020) have obtained single cell scale distribution of different glioma samples in a time section way. Even so, these data-based studies currently still face the difficulty of insufficient quality and quantity of data (Kirk, Babtie, and Stumpf 2015; Uthamacumaran 2022).

These data-driven studies are a continuation of the data explosion brought by the technological revolution witnessed in the past few decades. From the late 20th century to now, a vast amount of cancer-related knowledge was discovered by pioneers in cancer research, which was then summarized as a set of hierarchical hallmarks (Hanahan and Weinberg 2011) as a milestone. However, one of these pioneers asserted that cancer research is currently in false prosperity and urgently requires new theoretical foundations and mathematical tools (Weinberg 2014). The establishment of better theories and theory-experiment coupling, rather than solely relying on data that cannot generate true knowledge, was also called loudly in various fields of life science (MacArthur 2023; Smaldino 2019; Nurse 2021).

Such theories have long been budding. Wright (Wright 1932) and Waddington (Waddington 1957) introduced the metaphoric concept of adaptive/epigenetic landscape in evolution and development, respectively. Kauffman (Kauffman 1969) introduced Boolean dynamics to simulate gene regulatory processes, demonstrating the feasibility of using dynamics processes to model gene regulation and initiating the quantification of landscapes. Decades later, with the accumulation of necessary knowledge and data, the first real-data-based quantified landscape was obtained (X.-M. Zhu et al. 2004). After that, the Endogenous Network Theory (ENT) (Ao et al. 2008) was proposed, which argued that cancer and other complex diseases are endogenous states in the landscape emerging from the dynamics of the cellular-molecular network shaped by evolution, serving as a constitutive structure at the cellular-molecular level of the evolution dynamics (Ao 2005; Buchanan 2013), which then addressed several questions in cancer and development (Yuan et al. 2017). Lord *et al*. (Lord et al. 2019) provided experimental evidence connecting the theory of gene network dynamics and experimentally observed phenotypes. Studies in early brain development (Giacomantonio and Goodhill 2010; Goodhill 2018) as well as in hematopoietic system development (Olariu and Peterson 2019; Rockne et al. 2020) have also been implemented based on a similar systemic perspective.

Herein, we adapt the systemic perspective aforementioned to construct a core endogenous network for glioma based on experimental evidence. The energy landscape near glioma depicted by network dynamics enables us to elucidate the functional role of GSCs within the glioma hierarchy, in order to provide new perspectives for developing novel glioma therapies.

## Results

### Construction of the endogenous network for glioma

The existence and importance of gene regulatory networks no longer needs further debate, but how to obtain a dynamically workable and biologically rational network remains in controversy (Kim et al. 2023; Badia-i-Mompel et al. 2023; Prokop 2022; Uthamacumaran 2022). As the first step, we chose a bottom-up way based on causality knowledge rather than a top-down data driven way to construct the network.

The core function modules and signaling pathways of glioma have been already well summarized (J. Chen, McKay, and Parada 2012; Sedo and Mentlein 2014). We selected the following functional modules and signaling pathways to construct the core endogenous network for glioma (Fig. 1A): RTK pathway (Berezowska and Schlegel 2011; Annenkov 2014; Lemmon and Schlessinger 2010; Zeng, Cui, and Gao 2015; Newbern and Birchmeier 2010), NF-κB pathway (Perkins 2012; Taniguchi and Karin 2018; Sun 2017; Zhang, Lenardo, and Baltimore 2017), Ras pathway (Perkins 2012; Taniguchi and Karin 2018; Sun 2017; Zhang, Lenardo, and Baltimore 2017), Akt pathway (Manning and Toker 2017; Vanhaesebroeck, Stephens, and Hawkins 2012; Janku, Yap, and Meric-Bernstam 2018; Fruman and Rommel 2014), HIF pathway (Choudhry and Harris 2018; Gonzalez, Xie, and Jiang 2018; Nordgren and Tavassoli 2011; Rankin and Giaccia 2016; Eltzschig, Bratton, and Colgan 2014), p53 pathway (Agostini, Melino, and Bernassola 2018; Wade, Li, and Wahl 2013; Joerger and Fersht 2016; Kastenhuber and Lowe 2017; Hafner et al. 2019), cell cycle (Banfalvi 2011; Gérard and Goldbeter 2016; Otto and Sicinski 2017; Harashima, Dissmeyer, and Schnittger 2013; Bertoli, Skotheim, and De Bruin 2013), cellular senescence (Wiley and Campisi 2016; Kritsilis et al. 2018; Calcinotto et al. 2019; Gorgoulis et al. 2019; Muñoz-Espín and Serrano 2014), apoptosis (Singh, Letai, and Sarosiek 2019; Man and Kanneganti 2016; Spencer and Sorger 2011; Lavrik 2010; Hata, Engelman, and Faber 2015; Czabotar et al. 2014; Fricker et al. 2018; Zmasek and Godzik 2013) and glia differentiation (Kyrousi, Lygerou, and Taraviras 2017; Hidalgo and Logan 2017; Tchieu et al. 2019; K. S. Chen et al. 2017; Kamakura et al. 2004; Kang et al. 2012; Klum et al. 2018; Kuhlbrodt et al. 1998).

**Fig. 1.**
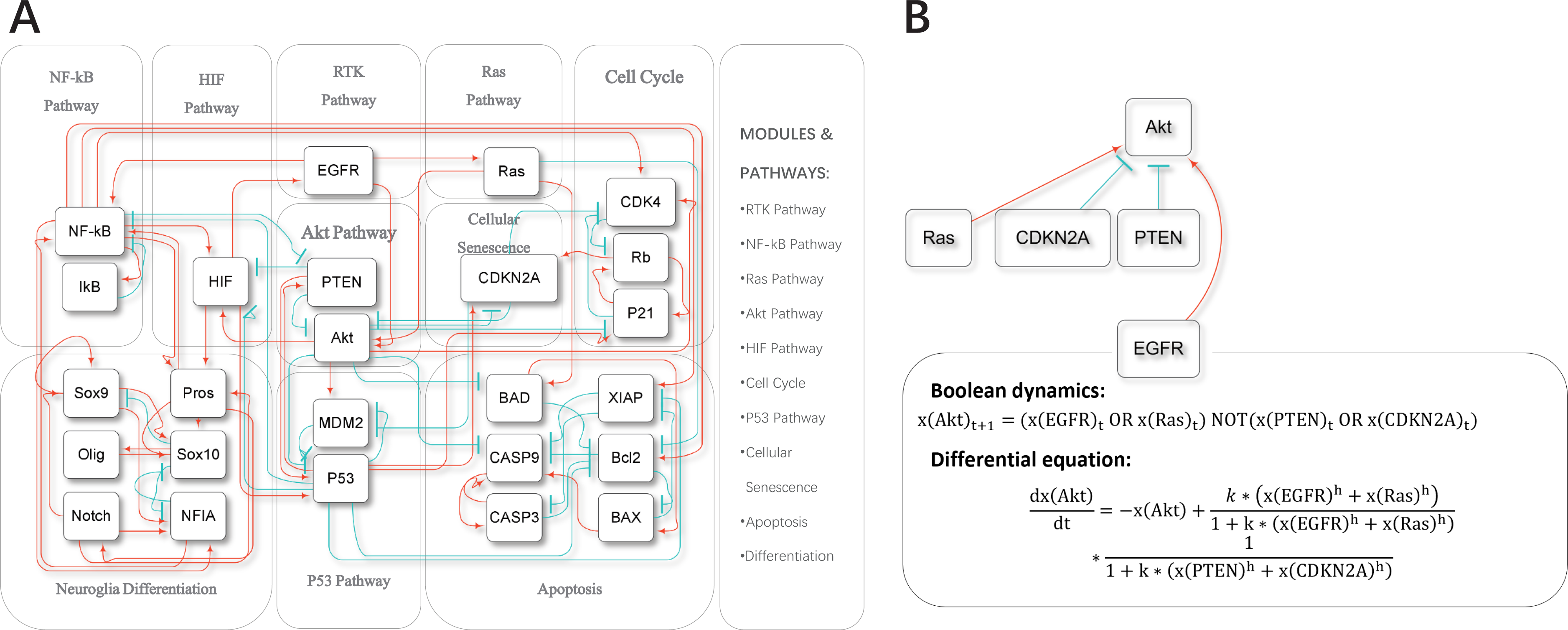
Endogenous network and network dynamics frameworks for glioma. **(A)** The core endogenous network for glioma. This network, constructed from experimentally driven causality knowledge, comprises 10 core functional modules or molecular pathways, including the RTK pathway, NF-κB pathway, Ras pathway, Akt pathway, HIF pathway, p53 pathway, cell cycle, cellular senescence, apoptosis, and glia differentiation. A closed a 25-node network with 75 edges formed by selecting key agents and interactions from modules and pathways, which includes 44 activatory interactions (indicated with red arrows) and 31 inhibitory interactions (indicated with blue T-shaped lines). **(B)** From the network to the dynamics. Two dynamic frameworks, Boolean dynamics and ordinary differential equations (ODE), were applied to this network. The dynamics equations for Akt are presented as an example. Here, x denotes the concentration/activity of core nodes, h represents the Hill coefficient, and k is the inverse of the apparent dissociation constant. A normalized degradation rate of 1 is used in this model, implying a degradation amount of -x per iteration. The full sets of equations are available in Supplementary Table 1.

The network consists of 10 function modules and pathways mentioned above, comprising 25 nodes and 75 edges, including 44 activatory edges and 31 inhibitory edges (Fig. 1A, Supplementary Table 1). For each node, we employed an equation incorporating both activatory and inhibitory effects to construct a 25-dimension equation set (Fig. 1B, Supplementary Table 1).

### Network dynamics results are robust under different computation frameworks

The Boolean dynamics was introduced into gene network dynamics by Kauffman in the 1960s (Kauffman 1969). As a discrete method, Boolean dynamics can quickly capture the overall structure. In this work, we used threshold functions equivalent to Boolean algebra to perform calculations (Fig. 1B, Supplementary Table 1). A Boolean equation set consisting of 25 Boolean equations was obtained for the network constructed above (Supplementary Table 1). With 10^7 random initial vectors, we obtained 62 attractors, including 16 point attractors and 46 linear attractors, resulting in a total of 116 states after expanding the linear attractors (Fig. 2A, Supplementary Table 1). By increasing the number of random initial vectors to 10^8, we obtained the same number and types of attractors (Supplementary Table 1). Additionally, by exhaustively considering all initial values (2^25=33554432), we obtained the same number and types of attractors (Supplementary Table 1). These results indicate that we have obtained all attractors under the Boolean dynamics.

**Fig. 2.**
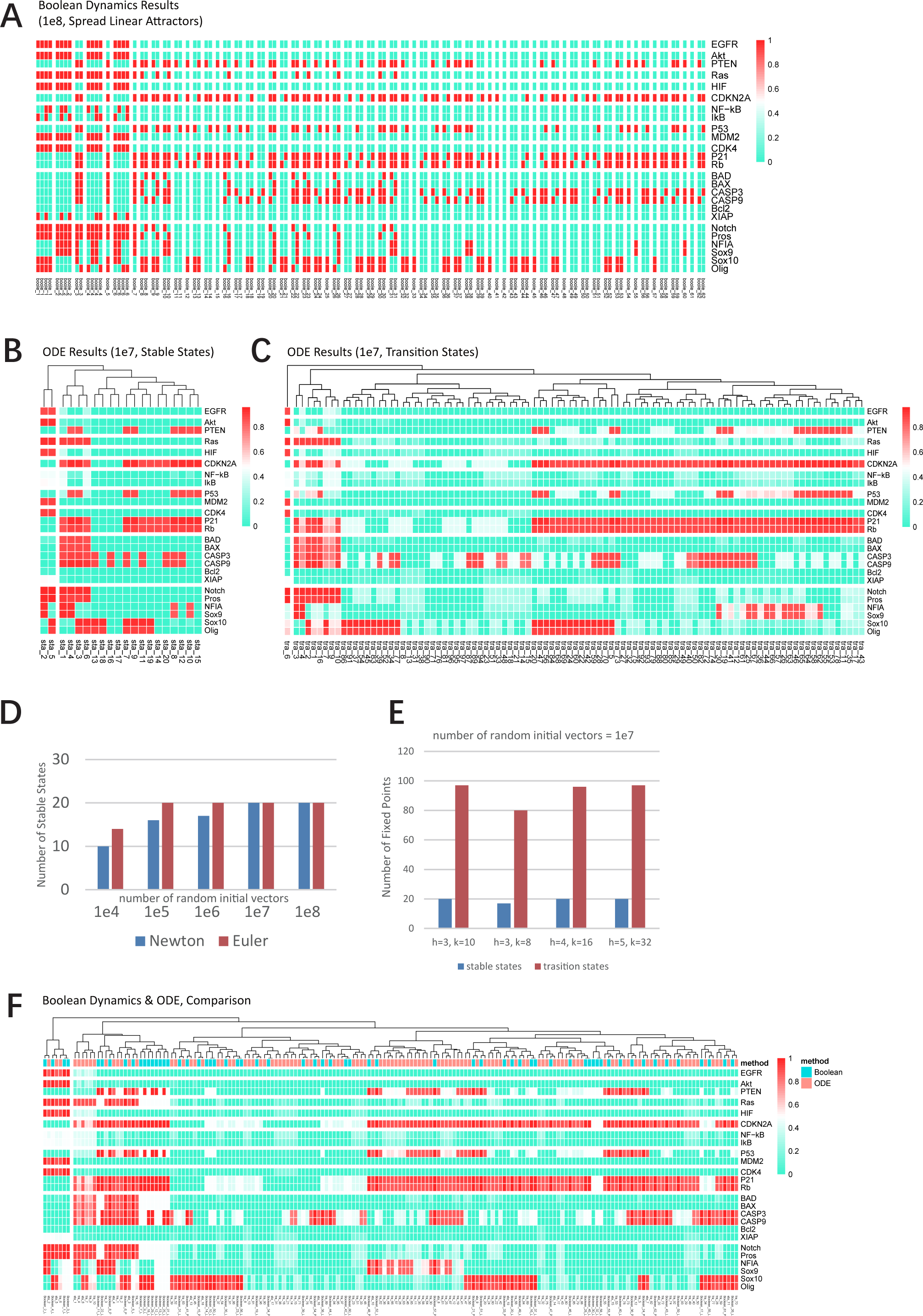
Dynamics results. **(A)** Attractor domains obtained using Boolean dynamics, with a random initial vector number of 1e8. States are represented with red for the active (1) and cyan for the inhibition (0). The x-axis corresponds to the identifier of each attractor domain, and the y-axis lists the core nodes. **(B) (C)** Fixed points obtained under the ODE framework. Fixed points, derived using ODE with a random initial vector number of 1e7, are hierarchically clustered by columns. Colors indicate the state, with red for active (1) and cyan for inhibition (0). The x-axis lists the identifier of each attractor domain, and the y-axis displays the core nodes. (B) Depicts stable states (stable fixed points). (C) Depicts transition states (unstable fixed points). **(D)** A comparison of the number of stable states obtained using different ODE algorithms. The count of stable states reached using Newton’s method (fsolve in MATLAB) and Euler’s method across various random initial vectors is plotted. The x-axis represents the count of random initial vectors, and the y-axis shows the number of stable states, with blue indicating results from Newton’s method and red from Euler’s method. Parameters used were h=3, k=10. **(E)** Comparison of the count of stable and transition states obtained under different parameters. The x-axis represents the parameters used, and the y-axis the counted number of fixed points. These results obtained under 1e7 random initial vectors and the Newton’s method. **(F)** Comparison between Boolean dynamics and ODE. After averaging the linear attractors from Boolean dynamics, they are hierarchically clustered by column alongside fixed points obtained from ODE. The x-axis denotes the identifier of the dynamic results, and the y-axis the core nodes. Grid colors indicate computation results, with red for active (1) and cyan for inhibition (0). Annotations above columns signify the computational framework, with red for ODE and cyan for Boolean dynamics.

While Boolean dynamics, as a discrete method, can quickly produce preliminary results, it fails to capture transitional states and simulate the consequence of small perturbations (Bornholdt 2008), therefore unable to determine the specific paths of transition between attractors. Experiments in biology often involve time-slicing, capturing features at the moment of death or detection, thus cannot guarantee that the system has reached equilibrium. Even if the system has reached a macroscopic equilibrium, noise-driven state transitions may occur at the microscopic level. Without transition states, it is impossible to determine how cell fate transitions occur. Stem cells cultured *in vitro* also exhibit characteristics of unstable equilibrium states that are prone to differentiation under small perturbations. Therefore, calculations that do not include transition states are incomplete. To address this issue, we need to use continuous computation frameworks.

The Langevin equation or stochastic differential equation (SDE) naturally corresponds to biological processes (Ao 2005). However, solving a n-dimensional nonlinear SDE, in this work n=25, would require excessive computation power, let alone there is currently no relevant knowledge for the random terms in this work. Moreover, the two commonly used stochastic integrals in applied mathematics and physics, Ito and Stratonovich, yield inconsistent results on the extreme points between SDE and ordinary differential equation (ODE) (Yuan, Tang, and Ao 2017). However, based on the mathematical tools developed previously, the extreme points of SDEs and ODEs are equal (Yuan, Tang, and Ao 2017). Therefore, solving a high-dimensional nonlinear SDE can be reduced to finding special solutions of high-dimensional nonlinear ODE, which are sufficient to depict the main features of the energy landscape of the system.

We chose the Hill equation for model construction in this work (Fig. 1B; Supplementary Table 1). Here, x represents the concentration/activity of core nodes, which is normalized to [0, 1]. The degradation rate is normalized to 1 in this model, so the degradation amount per iteration is -x. h and k represent the Hill coefficient and the inverse of the apparent dissociation constant, respectively. Since the equation has been already normalized, to ensure that the S part of the sigmoid curve is as close to the center as possible (i.e., f(x)≈0.5 when x=0.5), h and k need to satisfy the relationship k≈2^h. Based on previous experience, we chose h=3 and k=10 initially (Fig. 1B).

We obtained a set of 25 ODEs (Supplementary Table 1). We performed numerical calculations using uniformly distributed random initial vectors: generating a set of 25-dimensional random vectors corresponding to the concentration/activity of each node in the network as the starting point for each iteration. Both Newton’s and Euler’s method was parallelly applied to avoid bias.

We performed preliminary calculations with 10^4 random initial vectors. With Newton’s method, we obtained 10 stable states, and 14 with Euler’s method. To avert information loss due to insufficient sampling, we sequentially increased the number of random initial vectors tenfold until reached 10^8. The results converged rapidly on 10^5 random initial vectors to 20 stable states with Euler’s method compared with Newton’s method which the same 20 stable states were obtained on 10^7 random initial vectors. With 10^8 random initial vectors, Newton’s method and Euler’s method yield the same 20 stable states under the 10^7 condition (Fig. 2B, D; Supplementary Table 1). We also identified transition states similarly, the results converged to 97 transition states on 10^7 initial random vectors with Newton’s method and remained the same under 10^8 initial random vectors (Fig. 2C; Supplementary Table 1). Based on these results, it can be concluded that we have identified the vast majority, if not all, of the stable and transition states under ODE.

Furthermore, we explored the parameter space to ensure the parameter robustness. The results indicated the presence of bifurcations between h=3, k=8, and h=3, k=10, but 85% of stable states and 82% of transition states were already obtained under the h=3, k=8 parameter set. The quantity and types of fixed points obtained were consistent among h=4, 5, k=2^h, and h=3, k=10 (Fig. 2E; Supplementary Table 1). Considering that core nodes in gene regulatory networks of vertebrates often exhibit high interconnection (Gil-Gálvez et al. 2022), it can be speculated that significant cooperative effects exist in the core endogenous network of vertebrates. These results indicate that the network we constructed is robust against parameter alteration.

We obtained 116 solutions with spread linear attractors under Boolean dynamics; and 117 fixed points under ODE, yielding almost identical quantities. Hierarchical clustering and the Euclidean distances show that every attractor under Boolean dynamics corresponds to at least one fixed point in ODE (Fig. 2F; Supplementary Table 1). These results indicate that the network we constructed is robust against dynamics methods.

In conclusion, the dynamics results we obtained are acceptable enough for further analysis. Therefore, we will continue our work based on results obtained under the condition of h=3, k=10, 10^7 random initial vectors.

### Potential energy landscape of glia

In previous work, we discovered a universal construction method for energy functions (Kwon, Ao, and Thouless 2005). Subsequently, energy functions in various nonlinear systems were constructively obtained (Ma et al. 2014; Yuan et al. 2013; Y. C. Chen et al. 2020) and the existence of energy functions in nonlinear systems was mathematically proved in two dimension condition (Gan, Wang, and Ao 2021). Therefore, we believe that for the system studied in this work which can be taken as a set of equations of motion, an energy function and related energy landscape exists. For a hypothetical energy landscape demonstrated in Fig. 3 A, the fixed points in the equation of motion correspond to the extremal points (red dots “peak” as unstable fixed points/transition states and blue dots “bottom” as stable fixed points/stable states) in its energy landscape (Fig. 3 B). The topological connections between these stable and unstable fixed points are effectively enough to describe the primary features of its energy landscape (Fig. 3 C) which bypass the task of obtaining a global energy function of such a high-dimensional nonlinear system (Yuan, Tang, and Ao 2017; Yuan et al. 2017).

**Fig. 3.**
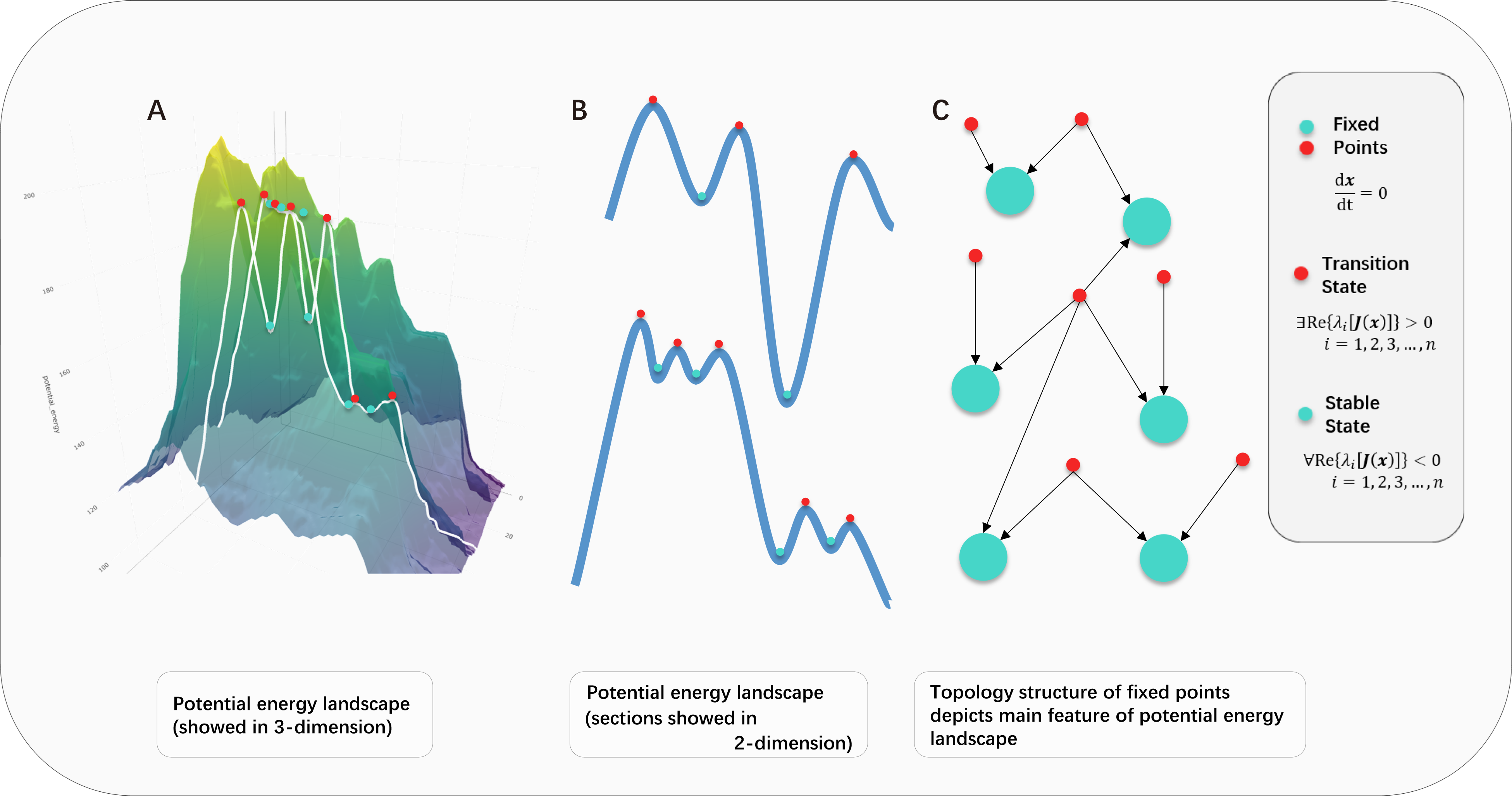
Schematic illustration: Using the topology structure of fixed points to depict the potential energy landscape. **(A)** A three-dimensional potential energy landscape schematic. The x-y plane represents a two-dimensional phase space, and the z-axis is potential energy. The “mountainous” surface depicts the potential energy surface, akin to Waddington’s epigenetic landscape used to describe development. White curves mark intersections between two orthogonal planes and the potential energy surface. Small dots on the curves represent fixed points, which will be further exemplified. **(B)** A two-dimensional projection of the potential energy landscape. White curves from (A) are mapped onto (B). (UP) indicates mapping of the y-z plane curve, while (DOWN) corresponds to the x-z plane curve. Small dots represent fixed points, corresponding one-to-one with those in (A). **(C)** A topological network mapping of the potential energy landscape. Small dots denote fixed points, correlating one-to-one with those in (A and B). Arrows point from higher energy fixed points towards lower energy ones. Red dots and cyan dots both signify fixed points, with red for unstable fixed points—those having at least one positive real part in the eigenvalues of the Jacobian matrix—and cyan for stable fixed points—those with all negative eigenvalues.

Specifically, we applied 10,000 small random perturbations to each transition state obtained earlier to simulate the system’s evolution from transition states to stable states, thereby obtaining connections between transition states and stable states which describe the main features of the energy landscape. We transformed the results into a topological connection graph of fixed points, where larger nodes represent stable states, smaller nodes represent transition states, the shape of the nodes represents the level of the apoptosis module, the color of the nodes represents the differentiation type, and the color of the node labels represents the activation status of the cell cycle (Fig. 4). It should be emphasized that the position of each node does not represent the energy.

**Fig. 4.**
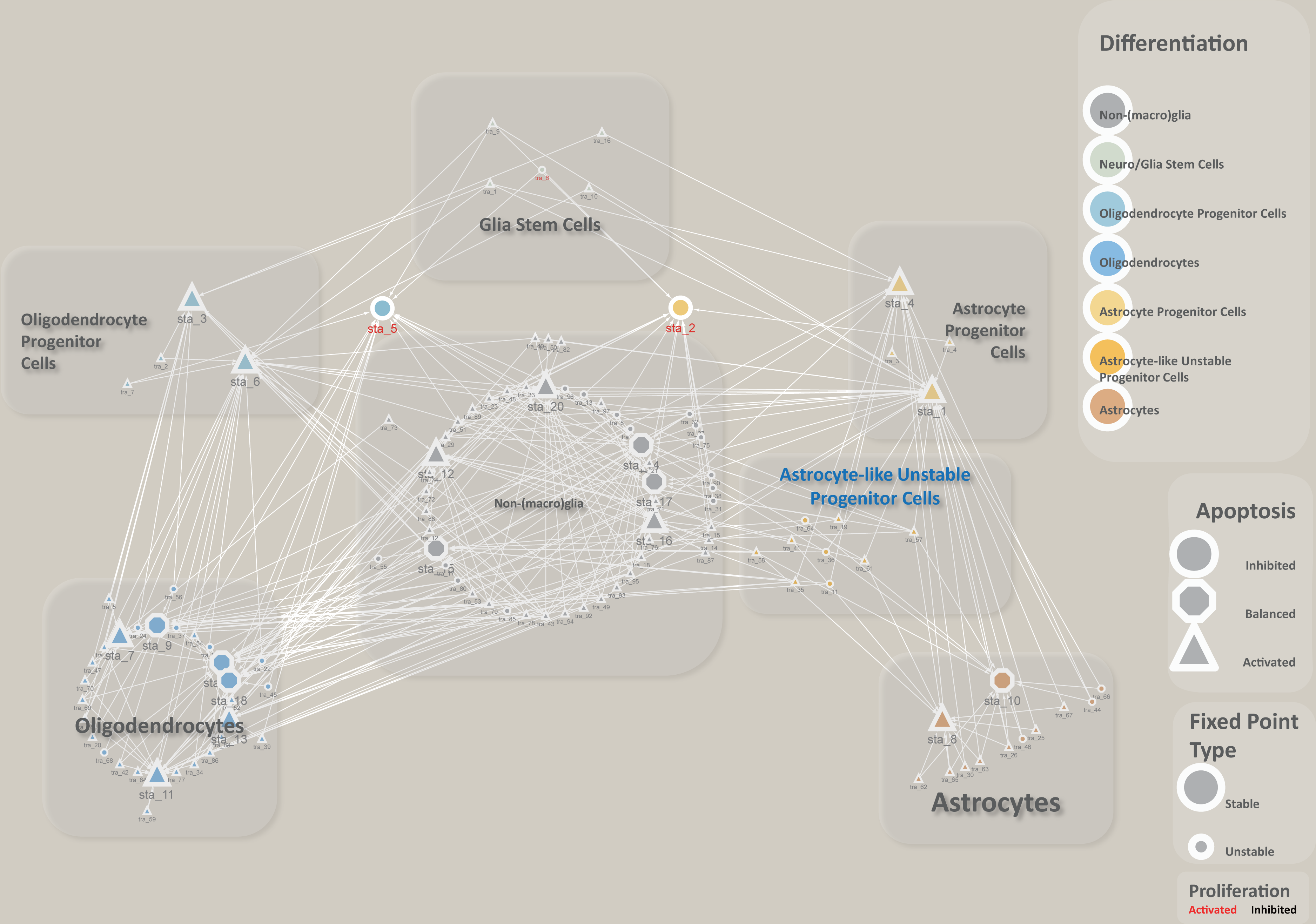
The global potential energy landscape of glia. The network topology of fixed points illustrates the global potential energy landscape for the glial system. Each node signifies a fixed point; large nodes indicate stable fixed points (stable states), and small nodes represent unstable fixed points (transition states). Node colors code for differentiation lineage, with yellow for astrocytes, blue for oligodendrocytes, and gray for glia stem cells or non-glia. Node shapes signal the apoptotic module state, with circles for apoptosis inhibition, octagons for balanced activation and inhibition of apoptosis, and triangles for apoptosis activation. Node labels in font color indicate the cell cycle module, with red for proliferation activation and black denoting proliferation inhibition. Arrows point in the direction of lower energy. Spatial relationships between nodes do not reflect relative energy levels.

Arranging the nodes according to their differentiation type, we obtained seven clusters of fixed points. One cluster of fixed points possesses high levels of stemness signatures, which consist of only transition states that will evolve into stable states with small perturbations, compliant with the characteristics of stem cells cultured in vitro, and was categorized as glia stem cells. Directly connected to this cluster of transition states are two clusters of fixed points, one possessed high level of both oligodendrocyte and stemness signatures and was categorized as oligodendrocyte precursor cells (OPCs), another possessed high level of both astrocyte and stemness signatures was categorized as of astrocyte precursor cells (APCs). A cluster of fixed points exhibiting low levels of all genes in the differentiation module was identified as non-(macro)glia. Two clusters of fixed points with the lowest levels of stemness and high levels of either astrocyte or oligodendrocyte differentiation signatures were identified as astrocytes or oligodendrocytes. Notably, there is a cluster of transition states that exhibits slightly high level of astrocytes-related signatures and slightly low level of stemness and oligodendrocytes-related signatures, which is capable of transitioning into multiple different differentiation types with small perturbations. We refer to this cluster as astrocyte-like unstable progenitor cells, which might correspond to NG2^+^ cells (Fig. 4).

After the arrangement, we found that the potential energy landscape of glia obtained encompasses the known glial differentiation hierarchy (Kriegstein and Alvarez-Buylla 2009; Laug, Glasgow, and Deneen 2018; Allen and Lyons 2018), indicating that the overall results up to this step under all mentioned modeling and computation are biologically acceptable. We then further explored the glioma-related implications based on these results.

### Locating the glioma states

To locate the glioma states, we validated the results at the module and pathway level at first. For a single-factor module or pathway, in this work, HIF, Ras, cellular senescence, and RTK, the value of the sole factor itself represents the level of the module or pathway. For a module or pathway that has both inhibitory and activatory nodes, with the hypothesis that each node in the core network as well as both the inhibitory and activatory part is equally important, we averaged the activation and inhibition levels and subtracted the inhibition level from the activation level, then normalizing the results to [0, 1] to obtain the module level results for multi-node modules. 0 represents complete inhibition, 1 represents complete activation, and values around 0.5 represent activation/inhibition balancing.

Stable states sta_2, sta_5, and transition state tra_6 exhibit high activation of the RTK, AKT, RAS, cell cycle, and HIF, as well as moderate activation of the NF-κB in module and pathway level. The p53, cellular senescence, and apoptosis modules are inhibited. This is fully consistent with the existing knowledge of glioma (J. Chen, McKay, and Parada 2012) (Fig. 5A, B). sta_2 exhibits a high level of Sox9, Notch, Pros, and NFIA, indicating a feature of low differentiated astrocytes; sta_5 exhibits a high level of Olig, Sox10, Notch, and Pros, indicating a feature of low differentiated oligodendrocytes; tra_6 exhibits a high level of Notch and Pros, as well as an intermediate level of Olig, Sox10, Sox9, and NFIA, indicating a feature of neuro/glia stem cells that have undergone gliogenic switch but have not yet differentiated into specific glial cell types (Elbaz and Popko 2019; Reiprich and Wegner 2015; Laug, Glasgow, and Deneen 2018; Wahlbuhl et al. 2012; Kato et al. 2011; Wang et al. 2020; Weider and Wegner 2017; Klum et al. 2018; J. S. Lee, Hoxha, and Song 2017; Glasgow et al. 2014; Samanta and Kessler 2004; Tchieu et al. 2019). Based on these results, we consider sta_2, sta_5, and tra_6 as glioma candidates, with sta_2 corresponding to astrocytoma cells, sta_5 corresponding to oligodendroglioma cells, and tra_6 corresponding to a type of glioma cell that embodied the feature of high stemness (Fig. 5A, B).

**Fig. 5.**
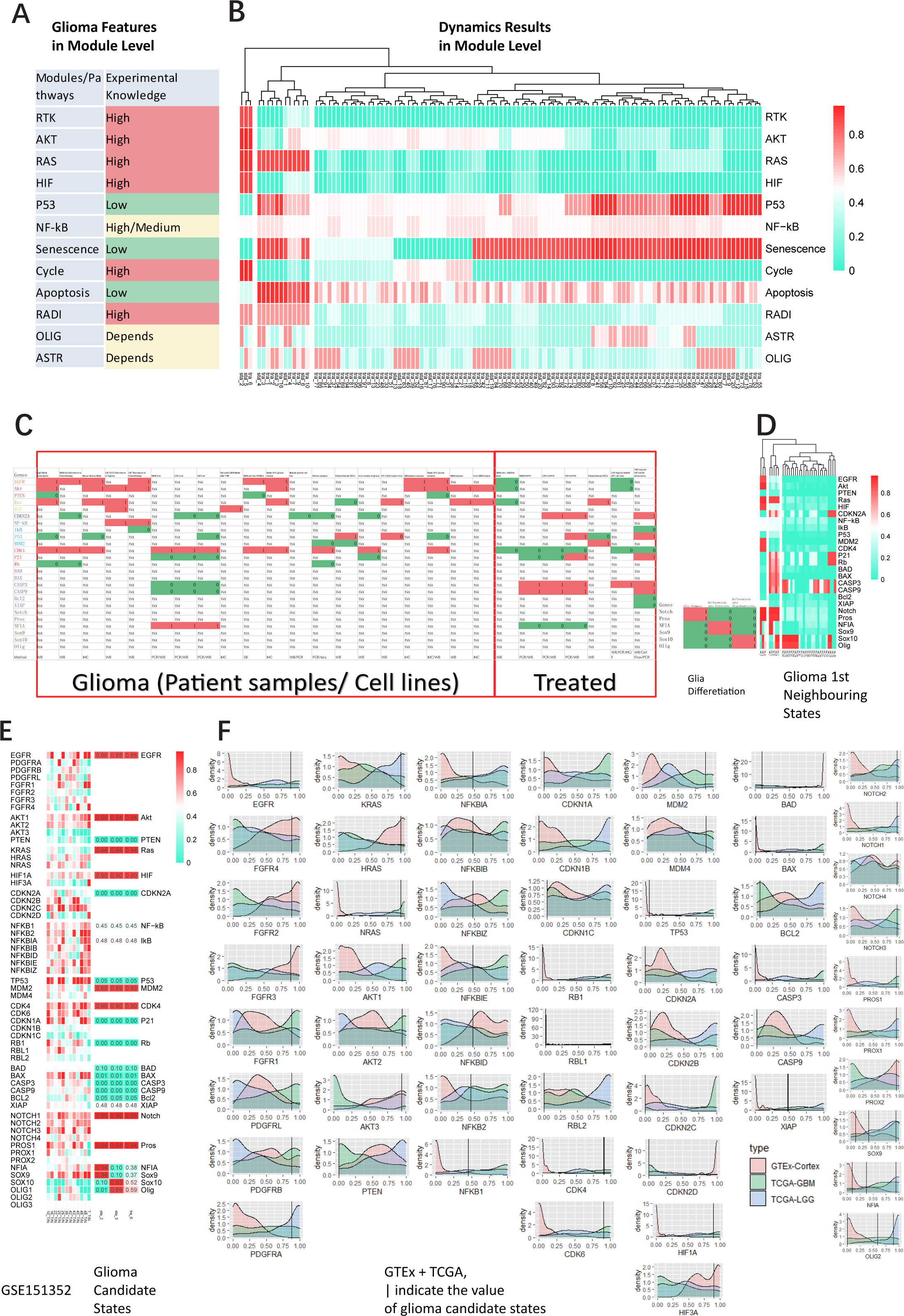
Comparisons between theory and experiment. **(A) (B)** Theoretical results compared with experimental data at the functional modules and signaling pathways level. (A) Knowledge about glioma summarized from the literature at the level of functional modules and pathways. (B) Theoretical calculations at the module level (ODE, Newton’s method, h=3, k=10, 1e7, same below). The far-left of the clustering tree displays glioma candidate states. **(C) (D)** Theoretical results compared with low-throughput data. (C) (LEFT) Core node-associated gene expression/activity data compiled from the literature, with columns representing individual reports. 1 denotes activation, 0 denotes inhibition, and na indicates the gene was not covered in that report. (BOTTOM-RIGHT) Typical states of transcription factors regulating glial cell differentiation summarized from the literature. (D) 1st Neighboring States of glioma candidate states as determined by theoretical calculations. **(E) (F)** Theoretical results compared with high-throughput experimental data. (E) (LEFT) Expression profiles of core node gene families from GSE151352, presented as fold changes between tumor and adjacent non-tumorous samples, normalized to the [0, 1]. (RIGHT) Theoretical calculation results for glioma candidate states. (F) Comparison of glioma candidate states with GTEx-TCGA expression profiles. TPM data from the cortex in GTEx and GBM and LGG samples in TCGA, with 150 randomly selected from each, normalized to the [0, 1] and plotted for the distribution of core node-associated gene expression. The x-axis represents normalized expression levels, and the y-axis shows distribution density; vertical lines indicate the level of that core node in glioma candidate states.

To make further confirmation of the correspondence of these three candidate states with glioma, considering that low-throughput methods, PCR, western blot, IHC, e.g., are still the gold standards for result verification in molecular biology, we then compared low-throughput experimental data with these three candidates at the molecular level. We obtained transcriptional and protein functional levels data on genes related to our model by reviewing literature from patient samples or cell lines(Chow et al. 2011; Cen et al. 2012; Marumoto et al. 2009; Jin et al. 2018; Tanaka et al. 2011; J. S. Lee et al. 2014; G. Lee et al. 2016; Thomas et al. 2001; Y. Zhu et al. 2005; Rollbrocker et al. 1996; Sato et al. 2011; Kfoury et al. 2018; Reilly et al. 2000; Guo et al. 2011; Zheng et al. 2008; Rajasekhar et al. 2003; Simon et al. 1999). After discretizing both our computational results and low-throughput data, we found that the consistency between the glioma states and the low-throughput data exceeded 95%, further confirming these candidates: sta_2, sta_5, and tra_6 as glioma states (Fig. 5C, D).

A single low-throughput experiment rarely covers all relevant genes in this model. Therefore, we further used high-throughput datasets to verify the computational results. We aligned RNA-seq datasets of LGG and GBM from TCGA with our candidate glioma states. Our computational results normalized the final activity of the factors, with 0 representing the lowest possible level of coarse-grained gene function in the system, and 1 representing the highest possible level. Since there are no normal brain tissues in this dataset, genes that may be highly expressed in LGG and GBM may not reflect their actual levels after normalization. Therefore, we jointly analyzed the TCGA data with the cortex RNA-seq dataset from GTEx. We selected genes related to the network in this work and performed double-normalization. After normalization, the glioma candidate states are organized into the same clustering branch with TCGA data rather than GTEx data after hierarchical clustering. We then made a more detailed comparison by using the distribution of every selected gene from the above datasets (Fig. 5 F). The comparison results for each module are as follows: In the RTK module, EGFR, FGFR1, and PDGFRA are consistent with the calculation results; In the RAS pathway, KRAS and NRAS are consistent with the calculation results; In the AKT pathway, AKT1 is consistent with the calculation results, while AKT2 and PTEN have multiple peaks in the experimental data, which are partially consistent with the calculation results; In the NF-κB pathway, NFKBIA, NFKBIE, NFKB1, and NFKB2 are consistent with the calculation results; In the cell cycle module, CDK4 and CDK6 are consistent with the calculation results, while the proliferation-suppressing genes CDKN1 (p21) and RB-related genes are completely opposite to our calculation results; In the P53 pathway, the experimental data of the classic “tumor suppressor gene” TP53 is completely opposite to our calculation results, while MDM2 and MDM4 are consistent with our calculation results; In the characterization of cell senescence, CDKN2 family genes, CDKN2A-C data all showed a double peak, the calculation results are only consistent with some sample data, only the expression data of CDKN2D is consistent with our calculation results; In the HIF pathway, HIF1A is completely consistent with the calculation results; In the apoptosis module, the expression of BAD and XIAP genes is consistent with our calculation results, BCL2 and CASP9 are partially consistent with our calculation results, and the expression of BAX and CASP3 is completely opposite to our calculation results; In the differentiation module, the results of NOTCH1-3, PROS1, PROX1 are consistent with the calculation results, SOX9 and NFIA data both have multiple peaks, each peak value can correspond to the calculation results, OLIG2 is extremely high in TCGA-LGG, and in TCGA-GBM is distributed uniformly, judged as partially consistent, SOX10 is not fully present among all datasets thus cannot be normalized nor been compared. We then integrated the comparison results to the core node level, for genes with consistent subtypes with core nodes, we set the comparison result to 1; for genes with opposite results, we set them to 0; for partially consistent results, we set them to 0.5; if the comparison cannot be made, they are not included in the statistics, and the average value is calculated for each module. The final average value of each module is used as the overall consistency rate, with a result of 76.5% (Fig. 5 F).

Since the above datasets came from different databases, to avoid bias caused by batch effects, we further used the dataset GSE151352 which consists of paired para-tumor and tumor tissue from GBM patients for comparison. We selected the core node-related genes and obtained the fold change of the relevant genes in each pair of para-tumor and tumor tissues, normalizing them to [0, 1]. Results from this comparison are not entirely consistent with the results obtained from TCGA-GTEx datasets. Among these: in the RTK module, EGFR and FGFR1 are consistent with the computational results; in the Akt pathway, AKT1, AKT2, and PTEN are consistent with the computational results; in the Ras pathway, HRAS and NRAS are consistent with the computational results; in the HIF pathway, HIF1A is consistent with the computational results; the marker of cell senescence, CDKN2A and CDKN2D in the CDKN2 family are consistent with the computational results; all subtypes in the NF-kB pathway are consistent with the computational results; the p53 pathway is opposite to the computational results; in the cell cycle module, all are consistent with the computational results except for CDKN1A; in the apoptosis module, all are inconsistent with the computational results except for BCL2 and XIAP; in the differentiation module, all NOTCH subtypes and PROS1 are consistent with the computational results, while NFIA, SOX9, SOX10, and Olig1-3 are partially consistent with the computational results. The overall consistency rate is 77.2% (Fig. 5 E).

Since the computational results in this work represent the functional level of the core nodes related gene, multi-omics analysis has already revealed that the correlation between the transcriptome and proteome is weak (Williams et al. 2016), further considering the inevitable cell type heterogeneity in tissue samples (Brat and Meir 2004; Suvà and Tirosh 2020), thus a perfect consistency between our results and the transcriptome cannot be expected. The consistency rate (76.5%, 77.2%) of high-throughput data validation is acceptable.

In summary, we can confidently assert that the glioma candidate states: sta_2, sta_5, and tra_6 correspond to glioma.

### Landscape near glioma

In the results aforementioned, we have portraited the energy landscape of the glial system and located the glioma states. To address the questions mentioned at the beginning, we need to take a close look at the landscape near glioma states. By selecting the 1st neighboring stable states which connect to the glioma states through only one transition state as well as the transition states connecting these stable states (Fig. 6 A, B), we obtained the energy landscape near glioma, consisting of a total of 12 stable states and 22 transition states (Fig. 6 C, D).

**Fig. 6.**
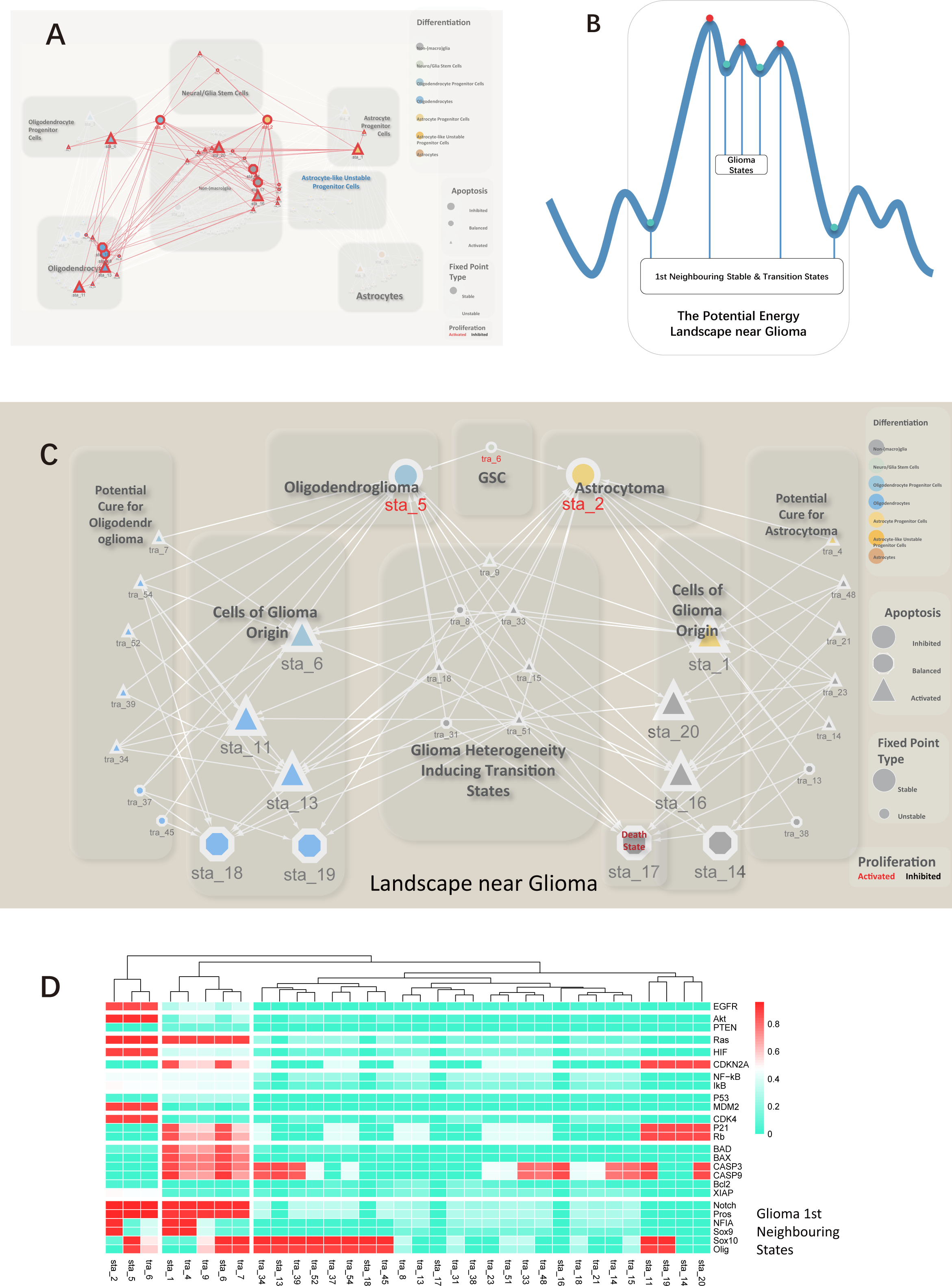
Landscape near glioma. **(A)** Locating the landscape near glioma states within the global landscape of the glial system. The transparent overlay represents the global landscape of the glial system, with the highlighted red network indicating the landscape near glioma states. **(B)** A two-dimensional representation of the 1st Neighboring States. Blue curves sketch the potential energy surface, with red and blue nodes corresponding to unstable or stable fixed points, respectively. Transition states connecting glioma stable states and those transition states connected to stable states constitute the landscape near glioma states. **(C)** The energy landscape near glioma. The 1st Neighboring States of glioma, rearranged according to differentiation type and connections to glioma states, yield the landscape near glioma states. **(D)** Profile of the energy landscape near glioma. Activities of core genes in all fixed points within the landscape near glioma states are hierarchically clustered by column. The x-axis lists fixed point identifiers, and the y-axis core genes. Red signifies active (1), and cyan inhibition (0).

### GSC state is the energy barrier separating astrocytoma and oligodendroglioma

All glioma states we acquired possess the feature of high stemness (Fig. 4; Fig. 5 C, D), which has been already observed by the analysis of single-cell sequencing data that most highly proliferative cells in samples of low grade glioma as well as glioblastoma exhibit high stemness (Suvà and Tirosh 2020).

Among these glioma states, a transition state, tra_6, is present. It possesses a signature of activated cell cycle, and high levels of glia stemness-related genes, Notch and Pros, as the mediated level of astrocyte differentiation-related genes, NFIA and Sox9, as well as oligodendrocyte differentiation-related genes, Olig and Sox10, representing low-differentiated glial cells that have undergone a gliogenic switch but remain high stemness (Laug, Glasgow, and Deneen 2018; Klum et al. 2018; Reiprich and Wegner 2015). tra_6 is an unstable fixed point, a transition state. With a small perturbation, it will and can only evolve towards the two glioma stable states, sta_2 and sta_5, consists with the knowledge of the functional features of GSCs (Lathia et al. 2015) (Fig. 7 A).

**Fig. 7.**
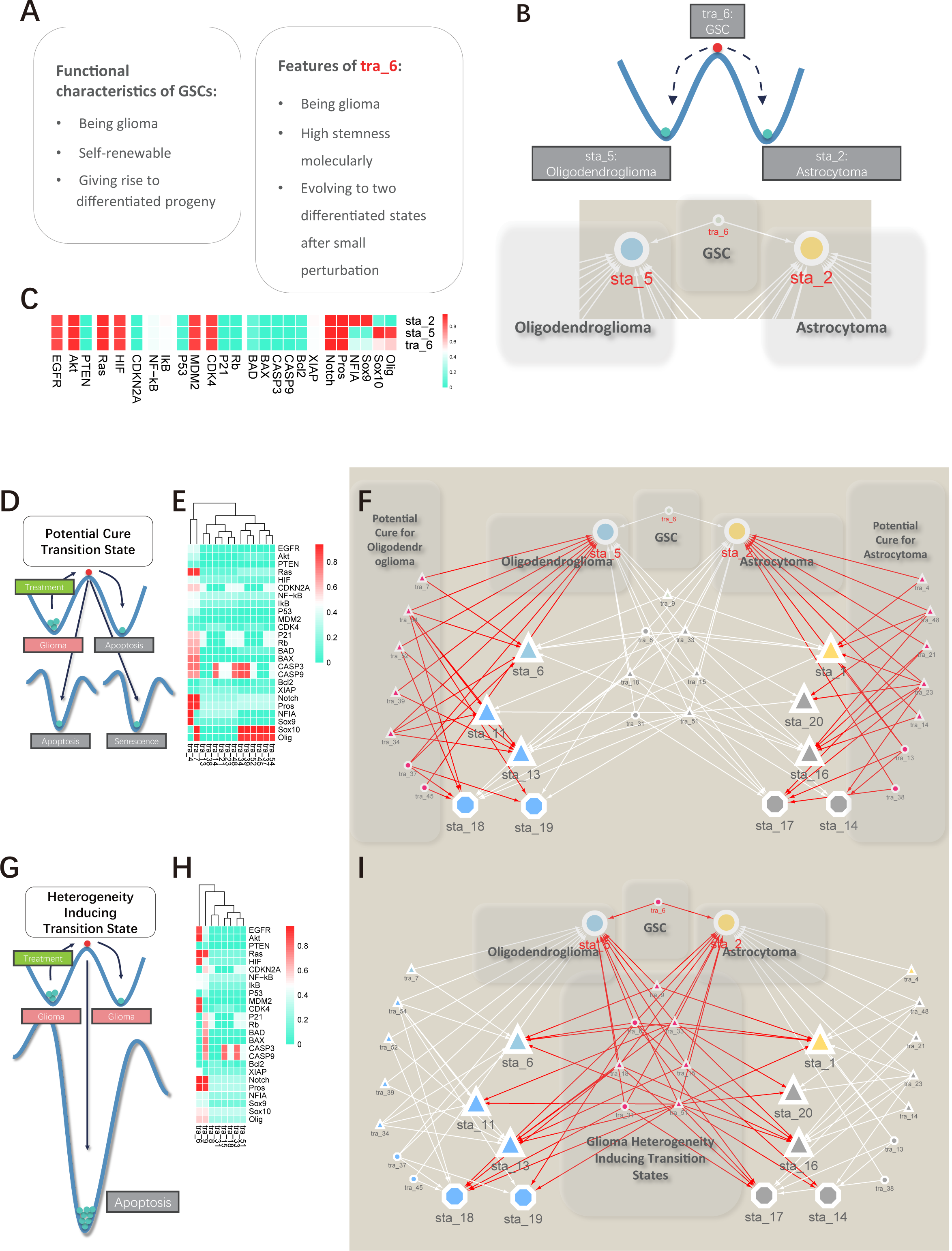
Transition states within the landscape near glioma states. **(A)** Theoretical versus experimental knowledge comparison for GSC states. **(B)** The GSC state is a transition state connecting exclusively to glioma stable states. (UP) Blue curves sketch the potential energy surface, with red and blue nodes corresponding to unstable or stable fixed points, respectively, same below. (DOWN) The connection of GCS state and glioma stable states. large nodes indicate stable fixed points (stable states), and small nodes represent unstable fixed points (transition states). Node colors code for differentiation lineage, with yellow for astrocytes, blue for oligodendrocytes, and gray for glia stem cells or non-glia. Node shapes signal the apoptotic module state, with circles for apoptosis inhibition, octagons for balanced activation and inhibition of apoptosis, and triangles for apoptosis activation. Node labels in font color indicate the cell cycle module, with red for proliferation activation and black denoting proliferation inhibition. Arrows point in the direction of lower energy. Spatial relationships between nodes do not reflect relative energy levels. Same below. **(C)** Theoretical calculation results of glioma states. The x-axis lists the core nodes, and the y-axis corresponds to the identifier of each attractor domain. Same below. **(D)** Schematic of potential cure transition states. **(E)** Profile of potential cure transition states. **(F)** Potential cure transition states in the landscape near glioma states. **(G)** Schematic of heterogeneity-inducing transition states. **(H)** Profile of heterogeneity-inducing transition states. **(I)** Heterogeneity-inducing transition states in the landscape near glioma states.

Further, the landscape indicates that a stable glioma state can evolve into another via tra_6 following significant targeted intervention or accumulated minor perturbations. When such transitions occur in large numbers, the transition state tra_6 can be clearly observed. This has also been observed experimentally, where a portion of rat GBM cell line C6 spontaneously exhibited stem-cell-like phenotype (De et al. 2020), as well as a developmental cell type “outer radial glia” being re-activated in GBM (Bhaduri et al. 2020).

A stable state needs sufficient time to evolve into another under the accumulation of minor perturbations, and the states that have crossed the energy barrier are exponentially distributed compared to non-transitioned states, meaning there are far fewer crossed cells than non-crossed ones. Suvà and Tirosh observed that the number of differentiated cells consistent with histological grading far exceeded those of another differentiated cell type (Suvà and Tirosh 2020), which again aligns with our theoretical results.

Based on the comparison between our theoretical results and the experimental observation mentioned above, only tra_6 in the entire glia landscape meets the features of glioma stem cells, thus we considered tra_6 to be the GSCs state (Fig. 7 B, C).

### Stable states near glioma are cells of glioma origin

In the energy landscape near glioma, all non-glioma stable states only need to traverse one transition state to transform into a glioma state (Fig. 6 A, B), meaning there is only one energy barrier to prevent these non-glioma stable states correlated cells from the fate of carcinogenesis, thus these states are all considered to be the most likely to transform into glioma states among the global energy landscape, in other words, cells correspond with these stable states are the cells of glioma origin, except the all-zero death state sta_17 (Fig. 6 C).

Sorted by differentiation module, they can be divided into groups of high stemness astrocytes or APCs (sta_1), high stemness oligodendrocytes OPCs (sta_6), differentiated oligodendrocytes (sta_11, sta_13, sta_18, and sta_19), and non-(macro) glia states that cannot be identified based on this study, which may be ependymal cells, neurons or mesothelial cells (sta_14, sta_16, sta_20) (Fig. 6 A, C, D).

60% of these stable states exhibit the feature of activated apoptosis module (sta_1, sta_6, sta_11, sta_13, sta_16, and sta_20), indicate the possibility that cells entered apoptosis program may also have the potential to enter the carcinogenesis progress (Fig. 6 C, D).

Previous researches reported that cellular senescence promotes glioma progression (Salam et al. 2023; Chojak et al. 2023), which also observed in our results. 60% of the glioma origin states (sta_1, sta_6, sta_11, sta_14, sta_19, and sta_20) exhibit activated cellular senescence module represented by the gene of CDKN2A which translates into p16^INK4^ (Fig. 6 C, D).

Experiments suggest that NSCs, mature astrocytes and OPCs could all potentially be cells of glioma origin (Zong, Verhaak, and Canolk 2012). Similarly, our results showed that astrocytes with high stemness and undergoing apoptosis are one type of cells of glioma origin, while differentiated astrocytes are not the cells of glioma origin. Oligodendrocytes with high stemness and undergoing apoptosis are also cells of glioma origin. Differentiated oligodendrocytes may also be cells of glioma origin, as well as non-(macro) glia cells which differentiation type are undistinguished based on this study solely.

Our results show that only tra_6 is the GSCs state in the landscape of glia, and tra_6, as a transition state, is unstable. That is, any small noise will cause it to evolve towards its neighboring stable states, in which only two glioma states exist (Fig. 7 B). Due to its unstable nature, the only condition of its long-term existence is always maintaining a subtle balance to stay in static equilibrium in the *in vivo* environment where noises constantly exist (Chang et al. 2008), otherwise it will soon transform into a stable state with less stemness. After that, it would only flash up during the path of the transformation from one stable glioma state to another. Thus, GSCs are unlikely to be the cells of glioma origin.

### Heterogeneity Inducing Transition States are Potential Mechanism Behind Therapy Resistance and Glioma Recurrence

In the energy landscape near glioma, all transition states (except tra_6) connect glioma origin states and glioma states, serving as potential routes for tumorigenesis and also as intermediate states between somatic glia and glioma (Fig. 6 C, D). These transition states can be divided into two categories based on their connections to glioma states:

The first type of transition states connects only one glioma state, such as those connecting only sta_2 and somatic states (tra_4, tra_13, tra_14, tra_21, tra_23, tra_38, tra_48) and those connecting only sta_5 and somatic states (tra_7, tra_34, tra_37, tra_39, tra_45, tra_52, tra_54) (Fig. 7 D, E, F);

The second type of transition states simultaneously connects both types of glioma states, including tra_6, tra8, tra9, tra15, tra18, tra31, tra33 and tra_51 (Fig. 7 G, H, I). Interestingly, all heterogeneity inducing transition states exhibit high stemness molecularly.

When driven from one glioma state through the first type of transition states, the system will only transfer into somatic stable states, which can be considered as potential cures (Fig. 6 C, LEFT & RIGHT; Fig. 7 D, E, F). However, if driven towards the second type of transition states, there is the possibility that the system transfers from one type of glioma stable state to another as well as multiple somatic states, resulting in tumor heterogeneity (Fig. 6 C, MIDDLE, Fig. 7 G, H, I). As long as the paths toward these heterogeneity inducing transition states exist, it would be difficult to reach a true cure, these treatment behaviors may further lead to an increase in the clinical grade of glioma.

### Conclusion

In this work, by taking a systemic theoretical perspective and integrating experiment-based knowledge, we constructed an endogenous network for glioma and depicted the energy landscape of the glioma system through network dynamics computation. By comparing theoretical computations with multi-level knowledge and data, we were able to locate the glioma states.

Our results revealed two stable states corresponding to astrocytoma and oligodendroglioma, interconnected by a transition state corresponding to GSC. This suggests that GSCs are unstable *in vivo* and are unlikely to be the cellular origin of glioma. The landscape near glioma states presented several stable states of diverse differentiation types, further confirming the multicellular origins of glioma theoretically.

In the landscape near glioma states, a group of transition states that simultaneously connected two types of glioma stable states displayed characteristics of non-proliferation and high stemness. If the system is driven through these transition states, this would lead to increased tumor heterogeneity. If these heterogeneity-inducing transition states are not blocked during treatment, the true cure of glioma will be difficult to achieve.

## Discussion

Differentiation therapy for glioma has already been called for (Laug, Glasgow, and Deneen, 2018; Suvà and Tirosh, 2020). Here, our theoretical results preliminarily suggest that the transitional states connecting two glioma stable states are responsible for glioma heterogeneity and resistance to radio-chemotherapy. These heterogeneity inducing transition states all demonstrate features of high stemness and low proliferation. Therefore, apoptosis-promoting therapies and differentiation-promoting therapies combined could be promising strategy in the treatment of glioma.

The model we used in this study is an “average field” model for the “average person,” and does not consider the genetic polymorphisms existing within populations. The network structure and parameters can be adjusted to further accommodate such genetic polymorphisms. Strategies for molecular therapies already target the core genes in our model (Rajesh et al., 2017). Our method can serve as an *in silico* navigation tool for developing personal cocktail therapies. However, we must concede that our 25-node model is overly coarse-grained, and future work should refine and enlarge the model.

Experimental validation of the theoretical results in this study is all retrospective. The following prospective experiments, such as investigating the presence of GSCs in the purified glioma cell line or implant tumors originated from differentially-purified glioma cells, can be used to verify our prediction that glioma stem cells serve as transition states connecting glioma stable states as well as that the heterogeneity and treatment resistance are mediated by heterogeneity inducing transition states:

For glioma cell lines or primary glioma cells, knock-in fluorescent protein markers following differentiation-related transcription factors such as the Sox family, NFIA, Olig, Pros, and Notch family mentioned in this work. Using flow cytometry to purify astrocytoma-like tumor cells as well as oligodendroglioma-like tumor cells. Observing the fluorescence signal during *in vitro* culturing and passaging. We expect to observe homogeneous fluorescence signals becoming heterogeneous during culture and passage with ground noise. Using microfluidic chips for long-term *in vitro* observation. We expect to observe the cell fate switching as observed in the system described by Lord et al. (Lord et al. 2019). Performing tumor implant experiments with purified cells. We expect to observe more LGG phenotypes than GBM phenotypes (implant tumor group); if low-dose apoptosis-inducing therapy is applied, the incidence of GBM phenotypes will increase (implant tumor + low dose treatment group); if glioma differentiation is promoted during low-dose apoptosis-inducing therapy, the incidence of GBM phenotypes will be lower than in the implant tumor + low-dose treatment group.

Research on gene regulatory networks has significantly advanced our understanding in recent decades. However, the integration of this vast array of static knowledge into a systemic framework that can yield dynamic insights has remained a challenge. In this study, we have attempted to bridge this gap by integrating experimental knowledge to construct a core endogenous network for glioma. By incorporating the concept of the landscape, rooted in biological principles and extensively applied in statistical physics, we have sought to generate novel insights. Our theoretical findings, while promising, represent a preliminary step that calls for further experimental validation. They offer a glimpse into the complex interplay between theoretical models and experimental research, a relationship characterized by mutual reinforcement and continuous dialogue. As we move forward, it is with cautious optimism that we envision our approach contributing to a deeper understanding of glioma biology and, ultimately, to the development of more effective therapeutic interventions.

## Methods

### Network dynamics

#### Boolean Dynamics

Boolean dynamics is a useful and relatively swift tool to obtain architectural information of the network dynamics results (Bornholdt 2008). The typical Boolean function is shown in figure 1 using Akt as an example: each variable has a discrete value 0 or 1 representing low/high level separately; t denotes the current state of the variables, t + 1 for the next state; NOT, AND, and OR are the usual logic operators. An equivalent formulation with threshold function is used in Matlab for our computation, details of the algorithms have been described in Yuan et al. 2017, the functions used in this work can be found in Supplementary Table 1.

#### ODE

We use two different algorithms to calculate stable states, details of the algorithms have been described in Yuan et al. 2017, the functions used in this work can be found in Supplementary Table 1.

#### Algorithms 1 (Euler’s method)

We initiate the procedure by generating a random starting point, denoted as the initial vector. Utilizing the dynamic system’s rules, we enter a loop of iterative calculations to progressively update the vector using a time step multiplication factor. The iterative process is designed to continue until we reach a predefined number of steps, ensuring the vector’s convergence to stability, which is ascertained through specific convergence criteria involving preset thresholds. Once a candidate for stability meets the convergence criteria, we classify and record it as a stable state.

#### Algorithms 2 (Newton’s method)

We employed a numerical method to solve the nonlinear equation representing the rate of change of the system variable over time. Similarly, our process began by initializing a random vector as a starting point for the iterative procedure. Using the fsolve function in MATLAB, we applied Newton’s method to locate the solution vector that zeroes the function, which signifies a potential stable state. To ascertain the stability of this solution, we evaluated the eigenvalues of the Jacobian matrix at the solution point, considering only those solutions as stable states where all eigenvalues have negative real parts, and those solutions where positive eigenvalues exist are considered as transition states. This procedure was systematically repeated to compile a comprehensive list of stable states.

### State interconnection analysis

Our algorithm is intended to generate trajectories leading from each unstable state to its corresponding stable states. To facilitate this, we introduced minor perturbations to the system via small, random vectors while it was positioned at an identified unstable state. This enabled us to trace the pathways from the unstable state to each linked stable state. The details can be seen in Supplementary table 1.

### Modelized the ODE results

In our study, the variable ‘x’ in the ordinary differential equations (ODE) represents the concentration or biological activity of nodes within the network. We have translated computational results to the module level to align with established biological knowledge. Following equation was used to determine the module status:

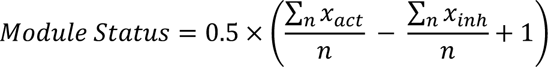

Nodes within functional modules and signaling pathways were categorized as either activatory (x_act_) or inhibitory (x_inh_), with the exception of the differentiation module. The final module status is determined by threshold values: an ‘OFF’ state is indicated when the Module Status is below 0.33, signifying inhibition; an ‘ON’ state is indicated when the Module Status exceeds 0.66, denoting activation; and a ‘Stable’ (STA) state is assigned when the Module Status falls between 0.33 and 0.66, reflecting a balance between activatory and inhibitory influences within the module. In a module with internal negative feedback, taking NF-κB pathway as an example: a STA state might suggest a balance similar to a 2:1 ratio of NF-κB subunits to their inhibitor pIkB forming a complex; an ‘ON’ state would imply a preponderance of free NF-κB subunits not complexed with pIkB and thus free to enter the nucleus and exert their function; an ‘OFF’ state would indicate a dominance of inhibitory genes, such as a high level of IkB relative to NF-κB subunits.

### High-Throughput Data Preprocessing

#### TCGA-GTEx

We acquired RNAseq data (TPM) from the GTEx cortex samples and the TCGA GBMLGG dataset. Randomly selecting 150 samples each from the cortex, GBM, and LGG collections, we performed a two-step scaling on the datasets. Initially, we computed the z-score for each sample to mitigate batch effects, and subsequently, we calculated the z-score for each gene. Values exceeding one standard deviation were capped to this threshold, followed by normalization to the [0, 1] range. The continuous distribution of gene expression was plotted by ggplot2 package in R.

#### GSE151352

Data from GSE151352 was retrieved from GEO database, converted into an exponential format (e), and its standard error was computed. We determined an effective information threshold for this dataset by averaging the standard errors of the genes with the top 200 smallest standard errors (≈30%). The original dataset was manipulated by subtracting the para-cancerous sample values from the tumor sample values to derive the fold change. A high expression threshold was set at ln(1.3), capping fold changes above this threshold to ln(1.3); a low expression threshold was set at ln(1/1.3), with values below this threshold adjusted to ln(1/1.3). Subsequently, the data was normalized to the range [0, 1].

